# A robust and automated algorithm that uses single-channel spike sorting to label multi-channel Neuropixels data

**DOI:** 10.1101/2020.12.19.423558

**Authors:** Zheng Zhang, Timothy G. Constandinou

## Abstract

This paper describes preliminary work towards an automated algorithm for labelling Neuropixel data that exploits the fact that adjacent recording sites are spatially oversampled. This is achieved by combining classical single channel spike sorting with spatial spike grouping, resulting in an improvement in both accuracy and robustness. This is additionally complemented by an automated method for channel selection that determines which channels contain high quality data. The algorithm has been applied to a freely accessible dataset, produced by Cortex Lab, UCL. This has been evaluated to have a accuracy of over 77% compared to a manually curated ground truth.

## I. Introduction

Neural electrophysiology (ePhys) has been a powerful tool that has enabled neuroscientists to study the brain over the past few decades. More recently, this has extended to the development of invasive Brain Machine Interfaces (BMIs) for medical and prosthetics applications [1]–[3]. Advances in integrated circuit and microsystems technologies [4]–[6] have enabled aggressive scaling in the number of recording sites that can be simultaneously observed [7]–[10]. This has enabled new exciting science, and more effective ‘high bandwidth’ BMIs, but has also exasperated the challenge with data throughput [11], [12]. There have been significant efforts towards developing methods for data analysis (e.g. spike detection and sorting) that compress the data whilst resolving the spatial information of the underlying neural activity. Computational methods have largely focused on accuracy [13]–[16], whereas hardware methods balance accuracy with real-time capability and power efficiency [12], [17]–[20].

Whether for analysis, or real-time decoding, the challenge however remains of how to assess performance in a real-world scenario without knowledge of the ground truth. There are generally 5 approaches here: (1) to use synthetic data where the ground truth is known [14]; (2) to simultaneously observe intracellular and extracellar activity thus knowing the single unit ground truth [13], [21]–[23]; (3) to establish the ground truth through manual labelling; (4) to estimate a hybrid ground truth using spatially oversampled extracellular recording [24], [25]; and (5) to use evaluation metrics that assess the algorithm without a ground truth [26]. Each of these methods have their strengths and limitations, e.g. for (1) it is challenging to establish realistic parameters such as signal-to-noise ratio, variability across channels, etc since these are highly electrode and tissue specific; (2) although definitively establishes the ground truth, this cannot scale beyond a few cells and can only be used acutely; (3) requires significant effort and is thus not scalable to large channel counts; (4) such data has only recently become available through high density active probes; and (5) avoids need for ground truth but is not widely accepted.

The work presented herein develops a novel automated algorithm for labelling Neuropixels data using the fact that neuropixels probes [8] can observe spatially oversampled data. In other words, the spacing between adjacent recording sites is significantly less than the distance single unit activity can be observed above background activity [8], [27]. One spike can therefore be observed across adjacent sites. By using this cross-channel information, ‘isolated’ spikes can be removed while any ‘missing’ spikes can be recovered. Therefore, the ground truth can be estimated with relatively good integrity. The remainder of this paper is organised as follows: Section II describes the datasets that are used and algorithm development; Section III describes how the labelling is evaluated, presenting results; and Section IV concludes this work.

## II. Material and Methods

### A. Test Dataset

This work uses a public dataset that was collected by Cortex Lab, University College London^1^, using Neuropixel probes (384 channels per shank, each sampled at 30 kHz) observing neural activity in the visual cortex [28]. This dataset has been sorted automatically using Kilosort [29] and then manually curated by Nick Steinmetz using Phy. This provides a reference dataset we compare our algorithm against in the absence of a genuine ground truth. The work presented in this paper uses a single short segment from this dataset (specifically the initial 33.3 s portion), with sample snippet shown in Fig. 1.

**Fig. 1.**
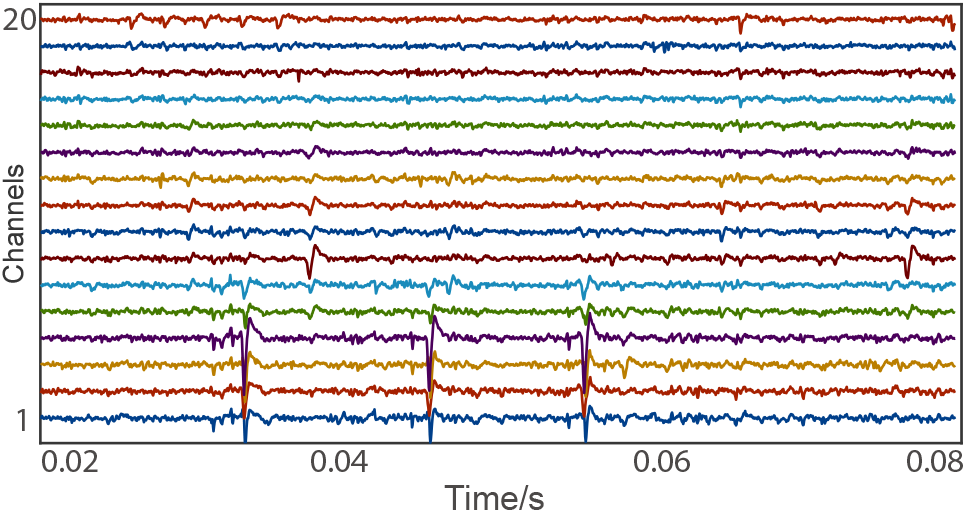
A sample of the raw data used in this work.

### B. Algorithm Development

The algorithm has been developed using MATLAB 9.5 (R2018b). This operates in four phases, as follows: (1) Initial labelling, to coarsely detect spike events across all channels with high sensitivity; (2) spike restoring and elimination, to recover missing spikes and remove false detected spikes; (3) spike clustering and merging, to separate spike events into those that originate from single unit activity (SUA) and multi-unit activity (MUA); and (4) to automatically select channels that contain good quality data, i.e. those that have been correctly labelled with high confidence. This process is illustrated in the flowchart in Fig. 2.

**Fig. 2.**
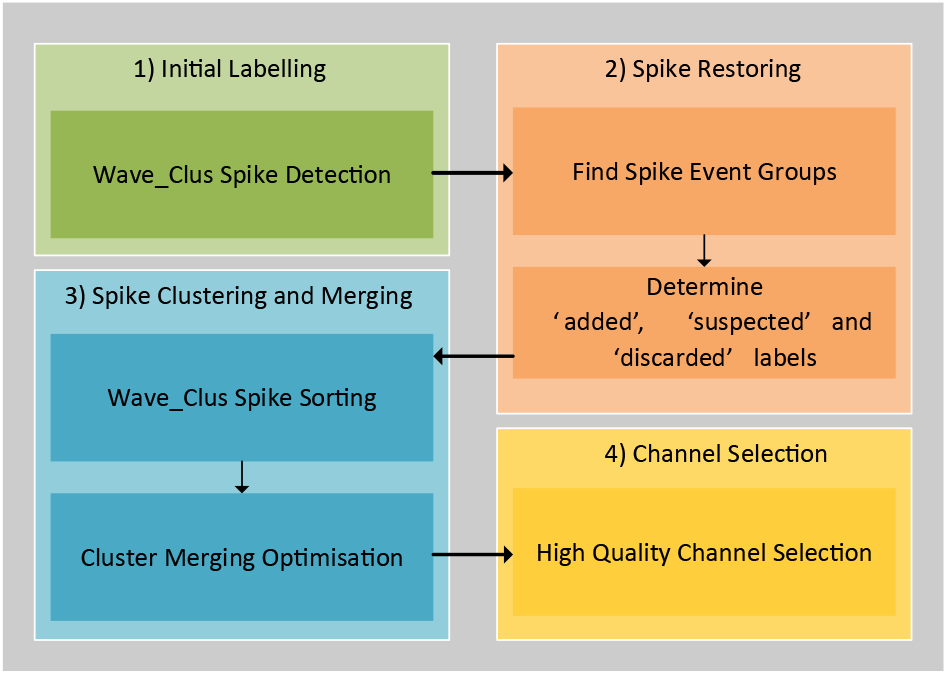
A flowchart of the proposed labelling algorithm.

#### 1) Initial labelling

The first step is to perform coarse (multi-unit) spike detection on each channel independently. This is achieved using Wave_Clus^2^, a widely used spike detection and sorting tool [14], [30]. This uses a global threshold for each channel to detect spikes, calculated using statistical properties as described by Eq. 1.

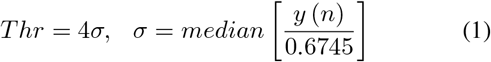

Wave_Clus intentionally operates its spike detection using a relatively low threshold such as to maintain a high sensitivity. This means that there are expected to be some false detections and also missing spikes as illustrated in Fig. 3(a).

**Fig. 3.**
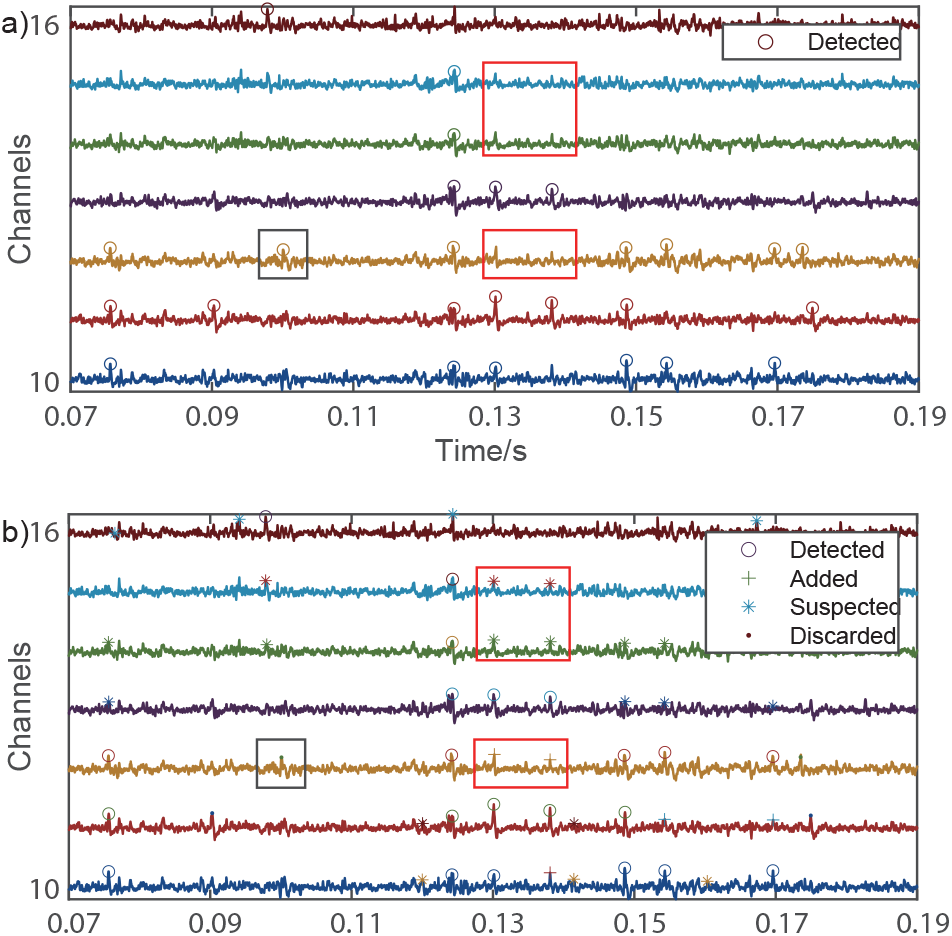
Example showing: (a) initial labels determined using MUA spike detection; (b) processed labels to include missing spikes, recovered based on spatial activity (red boxes), and discarded isolated spikes (black boxes).

#### 2) Spike Restoring

To compensate for any erroneous detections in the initial labelled (due to high sensitivity), we have applied two policies to recover missed spikes, and eliminate falsely detected spikes. This is made possible by the fact that Neuropixels provides spatially over-sampled data such that one spike can be observed across multiple spatial adjacent channels.

To utilise such cross-channel information, the spatial adjacency needs to be considered both horizontally and vertically. To locate the spatially adjacent channels, it is essential to understand the channel organisation across the Neuropixels probe shank. It can be observed (see Fig. 4), that the channels have been ordered in an appropriate way, i.e. spatially adjacent recording sites are also adjacent in channel number. The following rules are then applied:

- Spikes observed across closely adjacent channels that are detected within a 10-sample interval (±5 samples) are considered to be the same spike event. These are labelled using the ‘detected’ label.
- If at least 3 spikes are detected in a window of ±3-channels, then any undetected spikes are recovered, i.e. any gap is ‘padded’ with ‘added’ label. At the same timestep, two channels outer the channels that are closed to the window edge and have detected spikes, are labelled as ‘suspected’.
- if only 1 or 2 spikes are detected within this window, then they are assumed to be due to unwanted background activity and eliminated. These are labelled as ‘discarded’.
- This ±3-channel window is shifted in either direction until the boundaries of the spike event are determined.

**Fig. 4.**
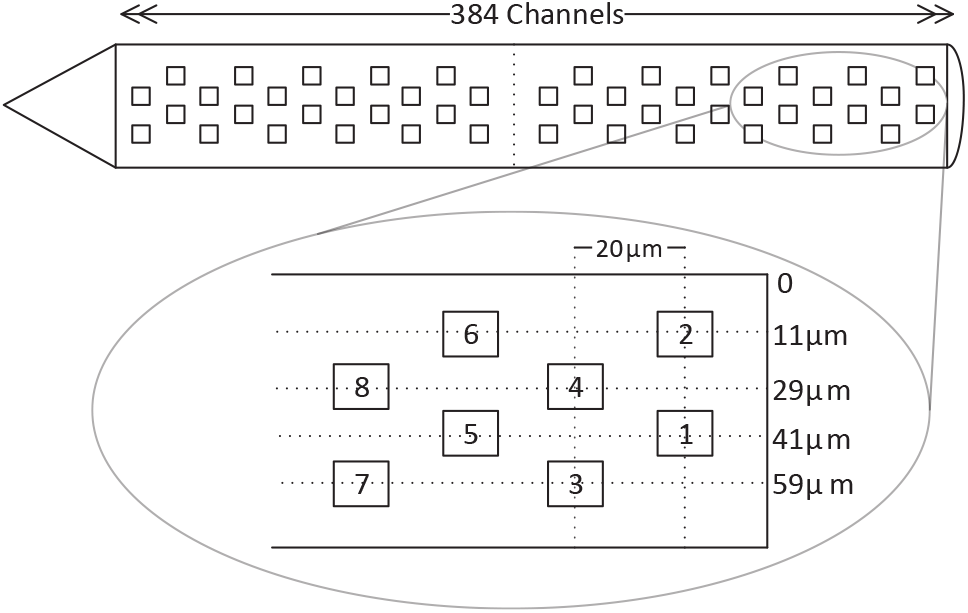
Spatial organisation of recording sites across the shank showing coordinates of the first eight channels.

Spikes are therefore labelled using either: detected, added, suspected and discarded. This labelling allows for some future adjustment of policies if needed. The different parameters (i.e. time interval, spatial window, etc) have been empirically determined based on the neuropixels dataset.

#### 3) Spike clustering and merging

As can be seen from Fig. 5(a), using simple spike detection can capture two ‘categories’ of spikes. These include single unit activity (SUA), those observed to have a larger amplitude (corresponding to neurons in close proximity to the recording site), and multi-unit activity (MUA), those observed to have a smaller amplitude (corresponding to background activity due to neurons further away from the recording site). Although this is sufficient for spike sorting (as the detection is proceeded by a classification step), an auto-labelling algorithm requires these spike types to be separated.

**Fig. 5.**
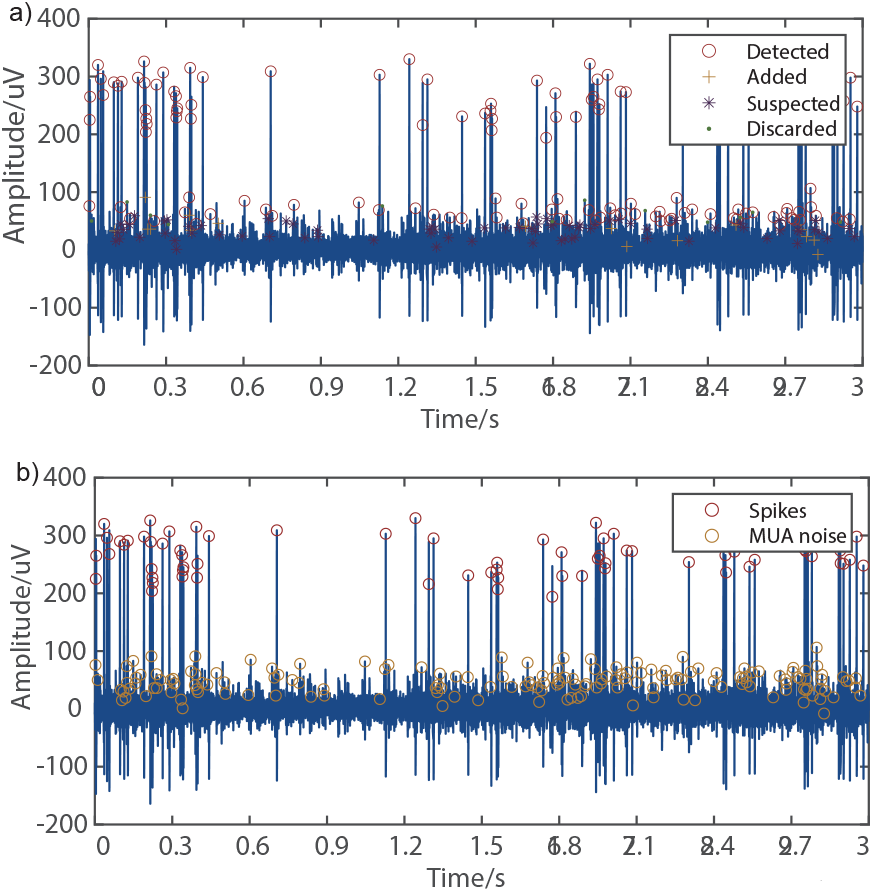
Example showing: (a) the processed labels; and (b) clustered labels, with SUA (valid spikes) and MUA (noise) separated into two groups.

One possible approach is to use a hard threshold to separate small amplitude spikes from larger ones. However, since these spike amplitudes vary from channel to channel, it is challenging to do so reliably without knowing the different spike classes (i.e. SUA).

We therefore adapt Wave_Clus spike sorting to achieve this. As the most distinguishable difference between the two types of spikes is their amplitude, we can apply spike sorting to determine the different clusters, and then merge the SUA clusters to separate these from the MUA cluster. This can be achieved by optimising the cluster merging for a given cost function – to maximize the difference between the average spike amplitudes of the two clusters. We do this optimisation using an exhaustive search – by trying all the combinations of how the spike sorted clusters can be merged into two groups. Additionally, in order to prevent small clusters disproportionately influencing the merging, we set a minimum limit on the size of a cluster of at least 5 spikes/sec. The two resulting spike categories after merging are shown in Fig. 5(b).

#### 4) Automated selection of high quality channels

The labels are now all tagged. However, there exist channels which are completely silent (thus containing no useful information), and other channels which are so noisy that no spikes can be reliably detected. This is a step that is normally done manually, and for this data set, 65 channels would be selected. However, to design a fully automated labelling algorithm, we have introduced a method to automatically select the higher quality channels. Firstly, all the ‘silent’ channels are discarded, in addition to channels with a signal-to-noise ratio (SNR) of under 13 dB. Of the remaining channels, the margin in spike amplitudes between the two spike categories (SUA and MUA) is then assessed, with only channels with a margin greater than 50 *μ*V are deemed ‘high quality’. In the dataset presented herein, 63 of the original 384 channels are selected. A sample taken from 16 of the selected channels is shown in Fig. 6.

**Fig. 6.**
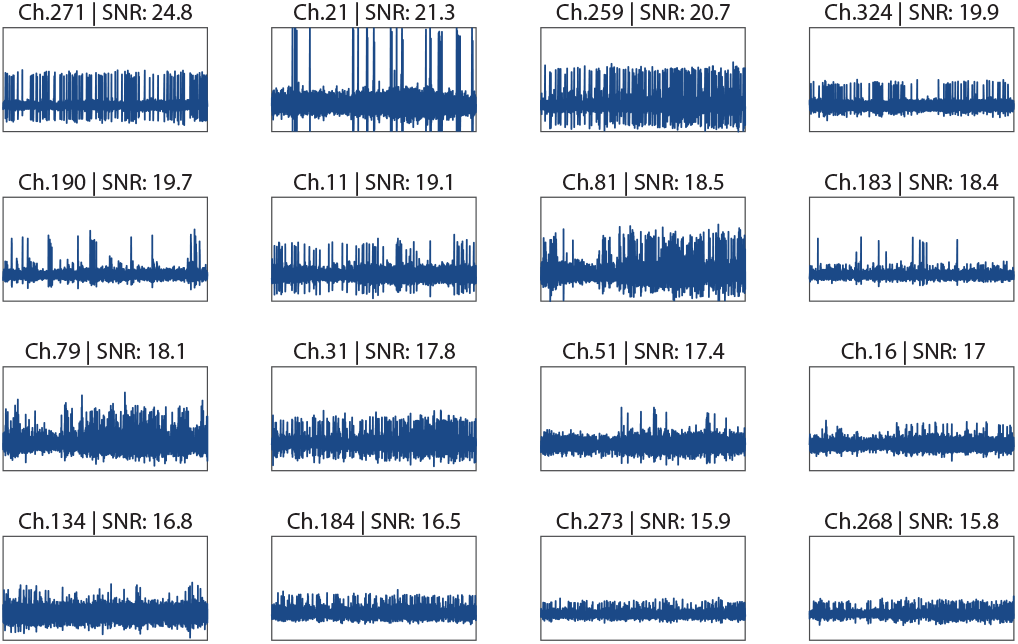
A sample of 16 channels (out of the 63 channels that are automatically selected) shown in descending order of SNR. Each waveform is annotated with channel number and SNR level.

## III. Results and Evaluation

The previous section has provided an automatically annotated dataset of 63 channels (65 if manually selected) with MUA noise and real target labels. In this section, we evaluate the effectiveness of the proposed algorithm in labelling the Neuropixels data, i.e. classification accuracy compared to the manually annotated labels. We also analyse the variability in the dataset we are using, to assess the sensitivity to different SNR levels.

### A. Labelling results

A summary of the detection results from the different stages, showing the assigned labels to the 33.3 s data segment is given in Table. I. ‘Events’ here refers to classified spike events, for example, a common spike observed across a group of channels.

**TABLE I.**
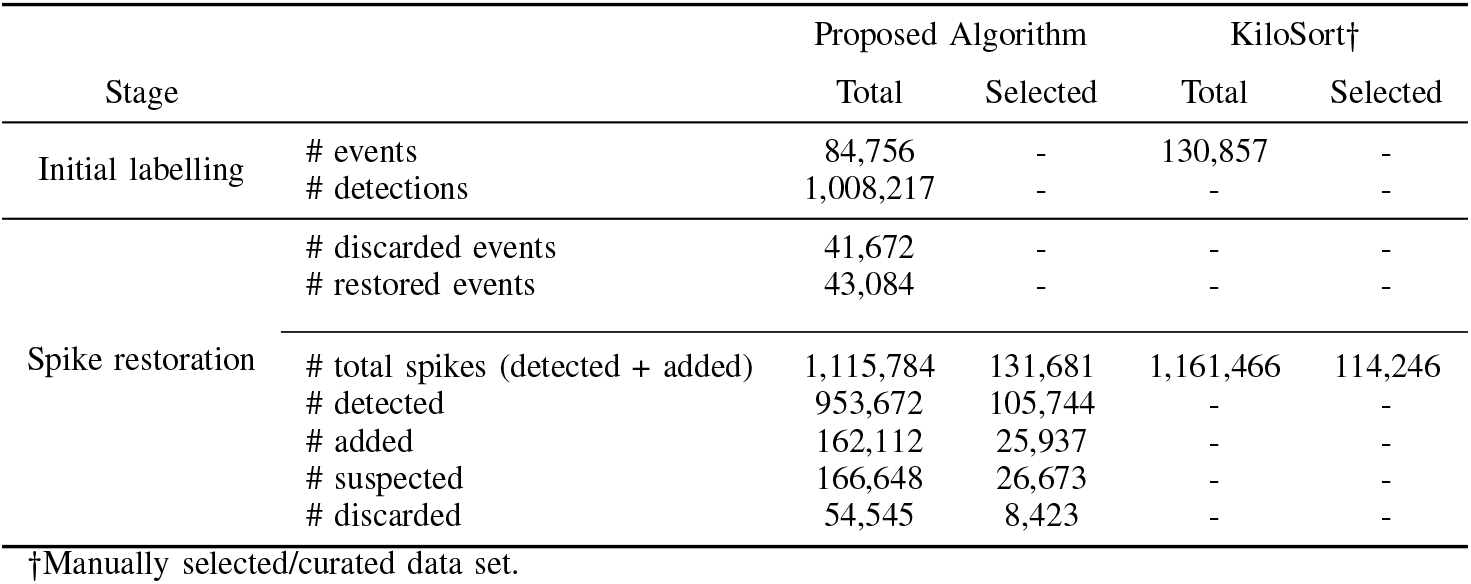
Summary of labelling results including the number of events and different labels.

### B. Algorithm Evaluation

#### 1) Spike Restoration

After spike restoration, the labelled spikes are shown in Fig. 3(b). Comparing to Fig. 3(a) some spikes are restored, as the red box shows, while some isolated spikes are discarded, as the black box shows. There are around 7,800 discarded spikes and 7,000 restored spikes across all channels every second.

#### 2) Spike Clustering and Merging

From Fig: 5(b), it can be observed that the spikes can be effectively separated into two clusters according to their amplitude. By manually checking across all channels, such an operation performs relatively well across 80% of the non-silent channels, which demonstrates the effectiveness of the spike clustering and merging method.

#### 3) Auto-selection

The auto-selection process that has been implemented identifies 63 channels, compared to 65 channels if manually selected. This is illustrated in Fig. 7. There are 42 channels in common, with a hit rate of over 87%. The auto-screening is therefore effective and accurate enough to replace channel manual selecting.

**Fig. 7.**
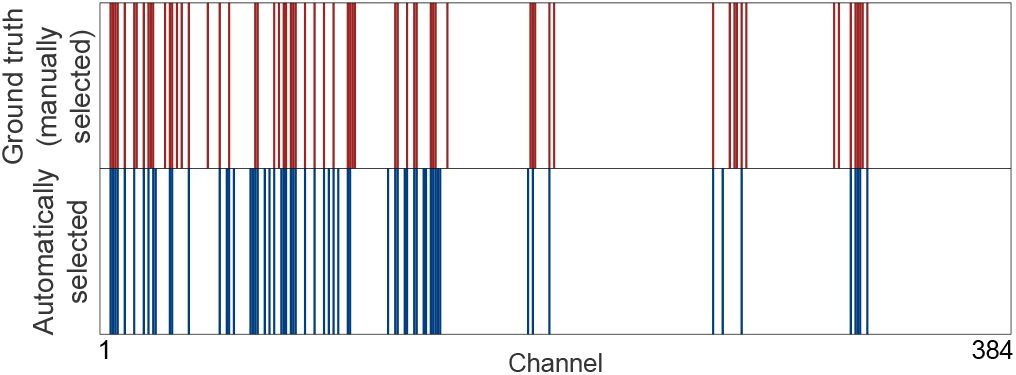
A comparison of manually and automatically-selected channels (with blue annotating channels that have not been selected).

### C. Dataset Evaluation

#### 1) Variability across different channels in dataset

From the plot in Fig. 6, the spike amplitudes, signal strength and noise levels can be observed to vary significantly across different channels. We have also assessed these features quantitatively. The range of spike peaks varies from the noise level (i.e. anything below 50 *μ*V) to over 300 *μ*V. This results in a SNR range from below 13.2 dB to 24.8 dB, with a mean value of 18.3 dB and standard deviation of 2.1 dB.

#### 2) Dataset Integrity

To evaluate the integrity of the algorithm, we calculate the similarity between the assigned labels (denoted as A) and the KiloSort results (denoted as *B*), which have previously been manually curated after sorting.

The detection accuracy we use is given by:

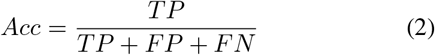

where TP stands for true positive, successfully detected spikes. FP stands for false positive, the locations where the signal is wrongly detected as spikes. FN stands for false negative, the undetected spikes.

We observe however that 10-20% of the spikes are assigned a channel number that has a slight offset from the channel that observes the highest amplitude. To address this, for all the channels that contain spikes that have been detected within the same time step, we pad labels in the 4 adjacent channels. This ensures that virtually all the ‘real’ spikes are correctly labelled but also introduce some undesired spikes, that degrade the accuracy we report below.

The SNR level and classification accuracy for each channel are first evaluated. These are used to determine the relationship between SNR and accuracy, shown in Fig. 8. The results show that the average accuracy (of the proposed algorithm) across the selected channels is 77% compared using the manually-labelled data as a ground truth.

**Fig. 8.**
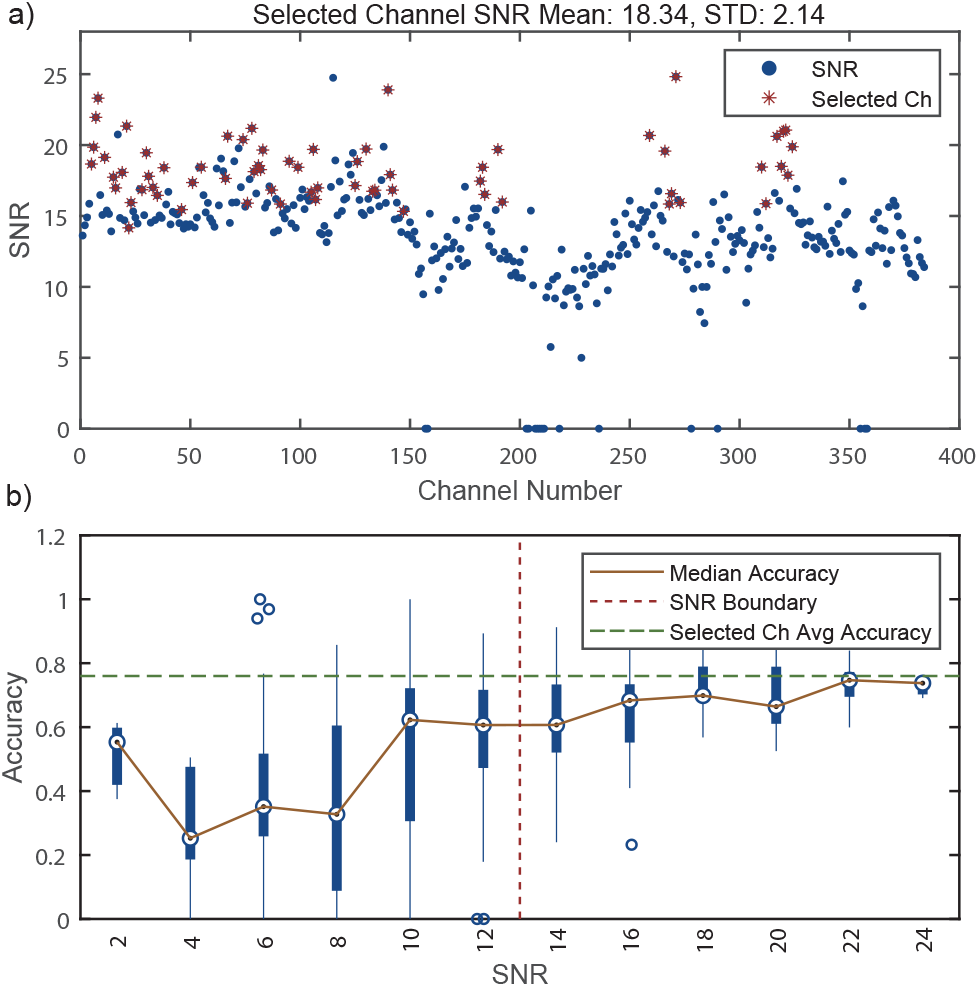
Variation and impact of Signal-to-Noise Ratio (SNR). Shown are: (a) variation of SNR across all channels; (b) classification accuracy across different SNR levels. The SNR boundary that corresponds to the average accuracy (0.77, across all channels) is 13 dB.

## IV. Conclusion

We have demonstrated a robust algorithm that can automatically label Neuropixel data that takes into account the spatial observation of spike signals. This provides a means for avoiding the need to manual label datasets to obtain a ground truth, or creating a synthetic dataset for evaluating spike processing methods. The labels that have been added to the dataset presented herein have been assessed to have an average similarity of 77% when compared to the manually-labelled dataset. ongoing work is further optimising the algorithm to improve similarity whilst increasing throughput, in addition to testing on other datasets. It is our intention to make both the algorithm and example labelled dataset available for wider use and further development.

## Footnotes

1 http://data.cortexlab.net

2 https://www2.le.ac.uk/centres/csn/software/wave-clus

